# Parameter fitting using time-scale analysis for vector-borne diseases with spatial dynamics

**DOI:** 10.1101/759308

**Authors:** Larissa M. Sartori, Marcone C. Pereira, Sergio M. Oliva

## Abstract

Vector-borne diseases are becoming increasingly widespread in a growing number of countries and it has the potential to invade new areas and habitats, either associated to changes in vectors habitats, human circulation or climate changes. From the dynamical point of view, the spatial-temporal interaction of models that try to adjust to such events are rich and challenging. The first challenges are to address the dynamics of the vectors (very fast and local) and the dynamics of humans (very heterogeneous and non-local). The objective of the present paper is to use the well-known Ross-Macdonald models, incorporating spatial movements, identifying different times scales and estimate in a suitable way the parameters. We will concentrate in a practical example, a simplified space model, and apply to Dengue’s spread in the state of Rio de Janeiro, Brazil.

## Introduction

Vector-borne diseases have now received increased attention because of their high potential of dissemination [1,2]. Mosquitoes of the species *Aedes Albopictus* and *Aedes Aegypti* are the most responsible for virus transmission, such as Dengue, Zika, Chikungunya and Yellow Fever. Due to lack of vaccination, basic sanitation, climate changes, and with increasing human mobility, such diseases are spreading and appearing in new regions, where the climate favors the proliferation of vectors. For instance, we may cite locations in Portugal, France and Italy where cases of Dengue and Chikungunya have already been recorded, and the United States with cases of dengue fever and Zika virus [3–6]. In addition, hosts may be infected in environments that are different from their place of residence because mosquitoes do not travel long distances, and this may lead to increased population heterogeneity and consequently in changes of the disease dynamics.

Dengue is currently the human viral disease with the highest number of cases, being an arbovirus of the family *Flaviviridae,* genus *Flavivirus,* is transmitted through the bite of female mosquitoes of the genus *Aedes* infected with the virus. It is estimated to be endemic in more than 100 countries, where climate favors the proliferation of vectors, and approximately half of the world’s population is at risk of contracting the disease [7–10]. For dengue, the current control measures are related to the vector population and its breeding sites, with the use of insecticides, adulticides and population awareness campaigns. More recently, control measures with *Wolbachia* bacteria have also been successfully tested, which prevents the vector from transmitting the virus [11,12]. Some vaccines have been tested and others are in testing phase, but the greatest difficulty is that such vaccines should be tetravalent, that is, they must be effective against the 4 existing serotypes of the disease [13,14].

Mathematical models used to describe indirectly transmitted infectious diseases have the interesting characteristic of coupling the dynamics of hosts and vectors, whose parameters have different time scales, the life cycle of mosquitoes is in days while humans life cycle is in years. Dengue is an example of infectious disease in which this occurs, in addition of having the particularity of different serotypes. Studies with spatial networks, or meta-populations, provide a way to understand the interactions between individuals in different scales, being a powerful tool to understand the characteristics of transmission in communities, regions and countries incorporating spatial heterogeneity [6,15–21]. In addition, as in modelling the dynamics of several vector-borne diseases, if the goal is to fit the model to real data, one has to deal with the asymptomatic cases, reliable data, in particular for the mosquitoes population, besides having to take into account the different time scales of vectors and hosts, which makes it difficult to study and understand disease dynamics [22–24].

In this work, we consider a basic compartmental model that divides the human population into susceptible S, infected I and recovered R (sometimes making it a little simpler), coupled with susceptible *S_m_* and infected *I_m_* mosquitoes. This model depends on parameters such as the mosquito mortality rate, the transmission rate and the total vectors population, which are very difficult to measure. From this model we do the time scale separation of humans and vectors. As a consequence, we have that the mosquitoes equations do not appear explicitly in the model, and the new parameters depend only on the total mosquitoes population. Finally, we incorporate spatial movements, considering mobility between cities two to two. We will adjust this model to dengue incidence data, whose cities were chosen based on [4]. Thereby, our main purpose is to show that it is possible to have a good fit of this reduced model in the beginning of the infection, as well as to estimate parameters related to mobility between two cities.

First of all, our approach will be deterministic, we will not take into account stochastic effects or incorporate the element of chance in the models. Secondly, being more precise, the goal is to consider the effects of the spatial dynamics into the Ross-Mcdonald models and use it to fit to the real data. This can be done either by a continuous space domain, which in turn will give us Partial Differential Equations, local or non-local, or consider discrete networks in space, which will provide system of ordinary differential equations ODE. There are advantages and disadvantages to both approaches. From the mathematical point of view, there are several theoretical challenges in the continuous model, in particular if non-local operators are considered, even if one proves that the second approach can be viewed as an approximation of the first and that the dynamics must somehow converge. The second approach can more easily be used to fit the real data, since they are always discrete in nature. Since, in this work, we are interested in concrete data and fit the dynamics, we will concentrate in the second model. For the continuous model, one can refer to [25–27].

The paper is organized as follows. First we set up the local dynamics, that will be used to describe the dynamic in each city, we also identify the small parameter that will be used. Next we set up the network dynamic, introducing a diffusion operator. With this two ingredients, for completeness, we show a formal expansion that reflect the general ODE singular perturbation results. Finally we can estimate our parameters using a network found to represent the initial spread of the disease in the State of Rio de Janeiro, Brazil and present our results.

## Methods and Materials

### Transmission model

Let us consider a model based on the classic Ross-Macdonald model. In our model, the total human population (*N_h_*) is divided in susceptible (*S*), infected (*I*) and recovered (*R*) and it is coupled with the compartments of susceptible *S_m_* and infected *I_m_* mosquitoes with total population given by *N_m_*, this model is named *SIRS_m_I_m_*. The interaction dynamics between the compartments is described through a system of ordinary differential equations (ODEs):

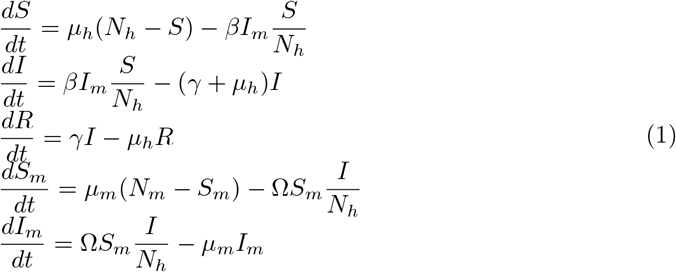

In this model, a susceptible human becomes infected at a transmission rate *β* when in effective contact with an infected mosquito, reciprocally, a susceptible mosquito becomes infected when it bites an infected individual at a transmission rate Ω. The parameter *μ_h_* represents the birth/mortality rate of humans, *γ* is the recovery rate of humans and *μ_m_* is the mosquitoes birth/mortality rate.

Assuming that birth and mortality rates are equal, we have that populations remain constant over time, that is, *N_h_*(*t*) = *S*(*t*) + *I*(*t*) + *R*(*t*) and *N_m_*(*t*) = *S_m_*(*t*) + *I_m_*(*t*), consequently we can easily obtain R(*t*) = *N_h_*(*t*) – *S*(*t*) – *I*(*t*) and *S_m_*(*t*) = *N_m_*(*t*) – *I_m_*(*t*) and then work with the equivalent reduced system:

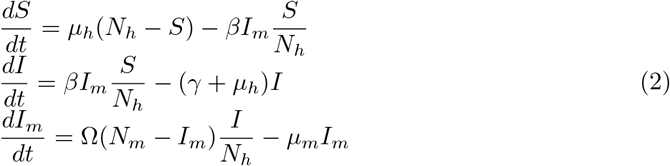

Considering that the life expectancy of an adult female mosquito is about 10 days [22], and a human life expectancy of 70 years, we have that the value of the parameter *μ_m_* = 7/10 (weeks) is bigger than the parameter of humans *μ_h_* = 7/(365 × 70) (weeks). Thus, to describe the time scales separation, we define 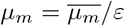, with 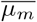 in the time scale of *μ_h_* [22,23]. So, setting up 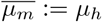, we obtain *ε* = *μ_h_*/*μ_m_* = 0.000392465 and 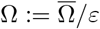, and then replacing these parameters in Model (2), we have

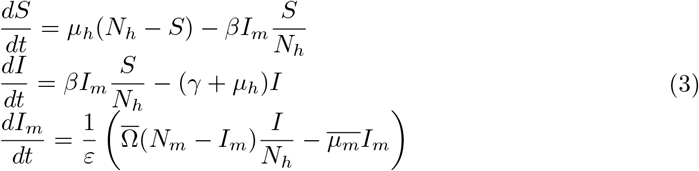

which is equivalent to

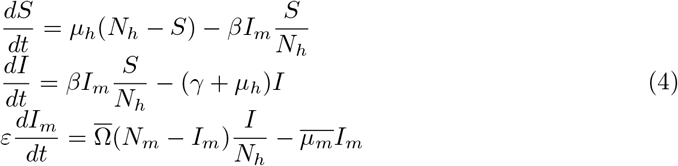

Making *ε* = 0 in the third equation, we have that *I_m_*(*t*) can be obtained as a function of *I*(*t*) at any time *t*:

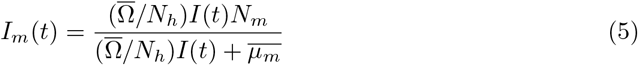

Replacing this value of *I_m_* Eq (5) into the equations of *S* and *I*, it results in

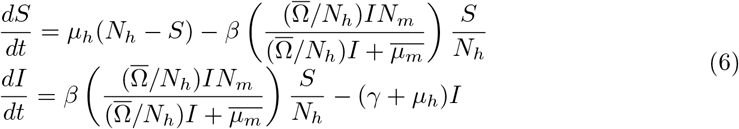

and then defining 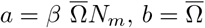 and 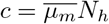, we have a new equivalent system without the variables related to mosquitoes:

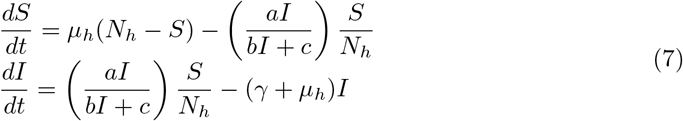

### Mobility

The *SIRS_m_I_m_* model characterizes the dynamics of a disease within a population. Thus, if we want to describe its transmission dynamics more realistically, we must consider a mobility network that includes interaction between populations. Such a network can be represented by a graph, where each node corresponds to a population and an arrow that leaves the population *r* to s means existence of mobility of individuals from population *r* to s [28].

Let *N_hr_* be the total human population that is registered in node *r* =1, 2,…, *M*, so that the disease dynamics in each location is described by the *S_r_I_r_R_r_S_mr_I_mr_* model. The parameter *d_rs_* ∈ [0,1] corresponds to the mobility rate from the population r to s per unit of time [18,28]. Accordingly, to include mobility in the Model (1), we must consider the inflow and outflow of humans in each compartment, and since mosquitoes do not move large distances, their respective compartments remain unchanged. Thus, the system of equations representing mobility between cities *r* and *s, r* ≈ *s*, is given by

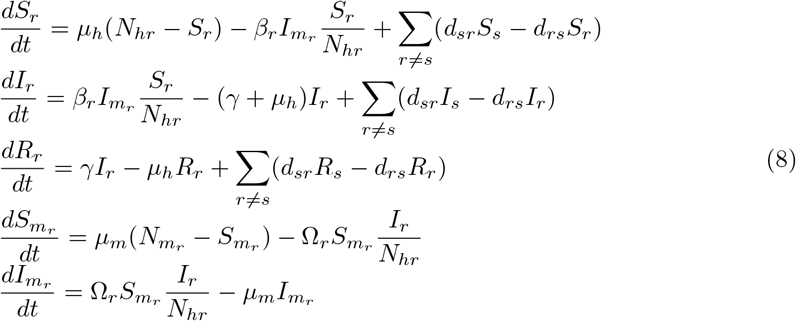

with initial conditions *S_r_* (0), *I_r_* (0), *R_r_* (0), *S_m_r__*(0), *I_m_r__*(0). Here, we suppose that the parameters are different for each location except the human birth/mortality rate *μ_h_*, and the human recovery rate *γ*. The total human population for each patch is given by *N_hr_* = *S_r_* + *I_r_* + *R_r_*, so

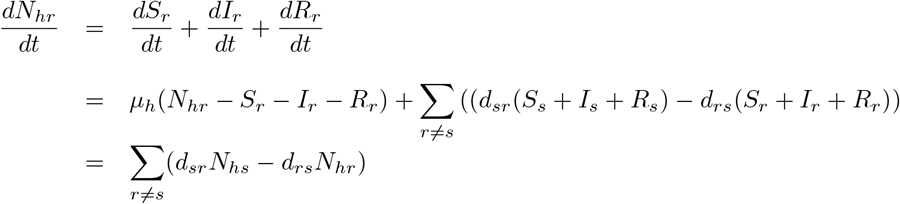

Similarly, we include mobility in (7), resulting in the ODE system, named *S_r_I_r_*:

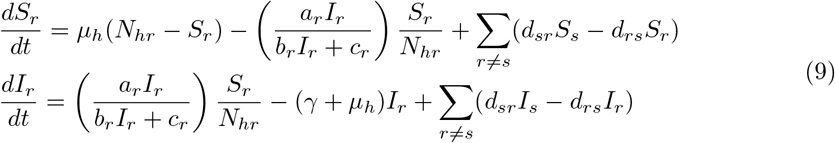

### Asymptotic expansion

Here we use a power series expansion to analyze the asymptotic behavior at *ε* = 0 of the perturbed System (3) with mobility given by (8). In order to do that, let **S, I** and **I_m_** be vectorial functions whose coordinates are denoted respectively by *S_r_, I_r_* and *I_mr_* for *r* =1, 2,…, *M*. We consider the following singular perturbed system of ODEs:

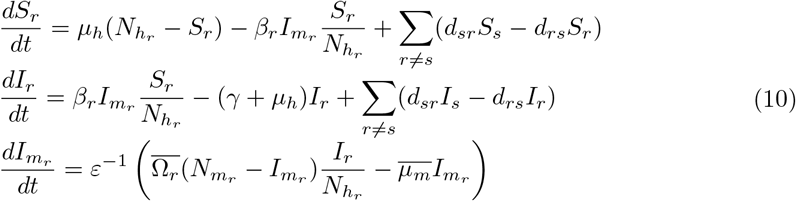

Now let us expand the solutions with respect to the small parameter *ε*, that is, let us assume that vectorial functions **S, I** and **I_m_** given by (10) satisfy

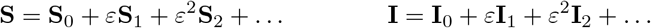

and

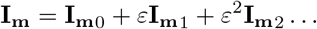

Thus, the time derivatives themselves set

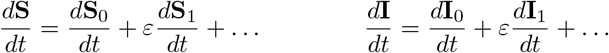

and

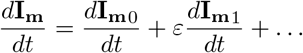

which gives us from the right-hand side of (10) that

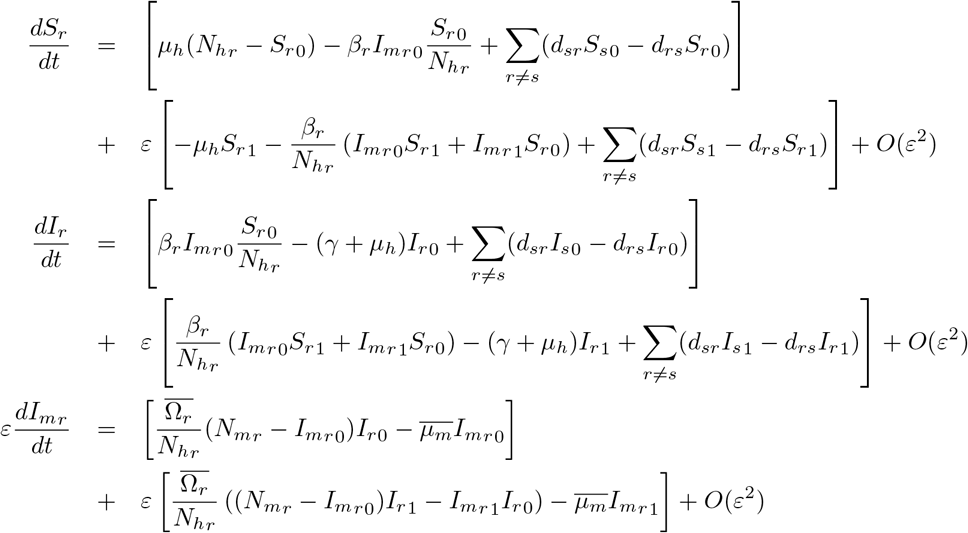

Then, if we plug these expressions in the System (10), we obtain at *ε* = 0 that

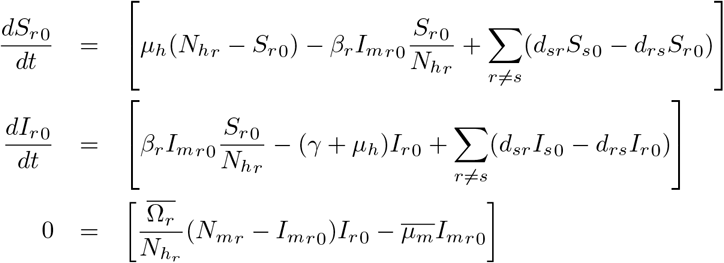

Hence, we get as in (5)

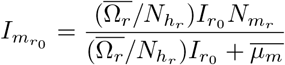

and then, we deduce the reduced equations

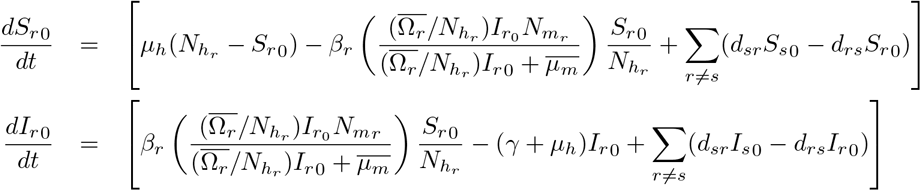

with initial condition *S*_*r*_0__ (0) and *I*_*r*_0__ (0). Thus, if we proceed as in (7) defining 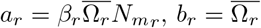 and 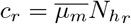, we can obtain the limit system (9) without the mosquitoes equations. Hence, the solutions **S** and **I** of the System (10) can be approximated to the solutions given by the *limit equation* (9). Indeed, under appropriated assumptions, it has been shown in [29][Theorem 4.4] (see also [22,23]) that the convergence is uniform in finite time with order *O*(*ε*) justifying our approach.

### Parameter estimation

We have the reduced System (9) which does not depend directly on the mosquitoes parameters. For the parameter’s estimation, we employed the **pomp** package implemented in language *R* [30], in order to obtain the values for: *a_r_, b_r_, c_r_*, *d_rs_*, 7 and the initial conditions of each location *S_r_*(0) and *I_r_*(0). The algorithm was applied to fit the *S_r_I_r_* model to dengue data of Brazilian cities which presents evidence of mobility according to the results obtained in [4].

The adjustment is done using the error of the least squares method. We set up a function that will calculate the sum of the squared errors which consists of the differences between the result obtained with the model and the data [30,32]. In our work we do the fitting of two time series each time. The model with time scale separation and considering mobility among two cities is given by:

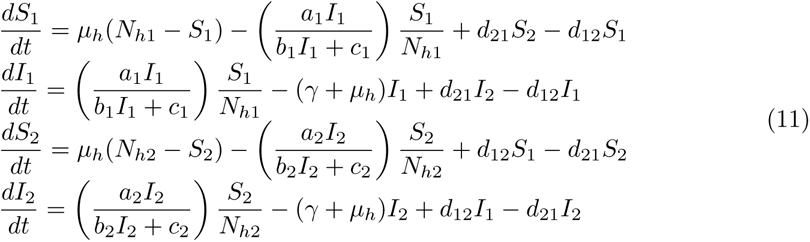

where

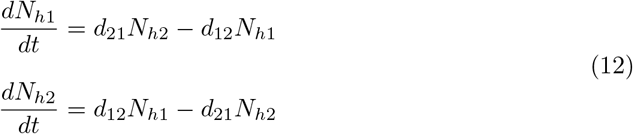

and the sum of the squared errors is:

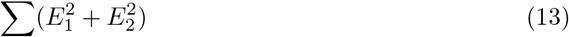

where *E*_1_ = *I*_1_ – *C*_1_, *E*_2_ = *I*_2_ – *C*_2_, and *C*_1_ and *C*_2_ are the weekly incidences of City 1 and City 2, respectively.

After set up the error function which is the objective function, we apply an optimization algorithm in order to minimize its value. To each parameter and variable can be given a lower and/or upper bound, then it is necessary to start with an initial value which satisfies the constraints and from this point, the optimization algorithm will search the parameter space for the value that minimizes the objective function. The algorithm uses function values and gradients to build up a picture of the surface to be optimized [30]. It is important to note that it is a ill-posed problem and a change in the initial conditions may change the estimated value of the parameters.

### Data

We will consider dengue data for our simulations, which were obtained from the brazilian Notification Disease Information System (SINAN) [31]. The year 2008 was chosen for results presentation due the high incidence of dengue cases in Rio de Janeiro state, considering the period from the 1st week to the 35th week of 2008. The number of reported cases per week (incidence per week) in Duque de Caxias, Itaboraí, Niterói and Nova Iguaçu are shown in Fig 1. These cities where chosen among all others based on [28], which used the ideas of [33, 34] to estimate an effective network that explained the epidemic in Rio de Janeiro. In this year, they detected that the disease started and spread to such cities, before spreading to the whole state. The fixed parameters are the total population of each city and the birth/mortality rate of humans (see Table 1), and the other parameters need to be estimated.

**Fig 1.**
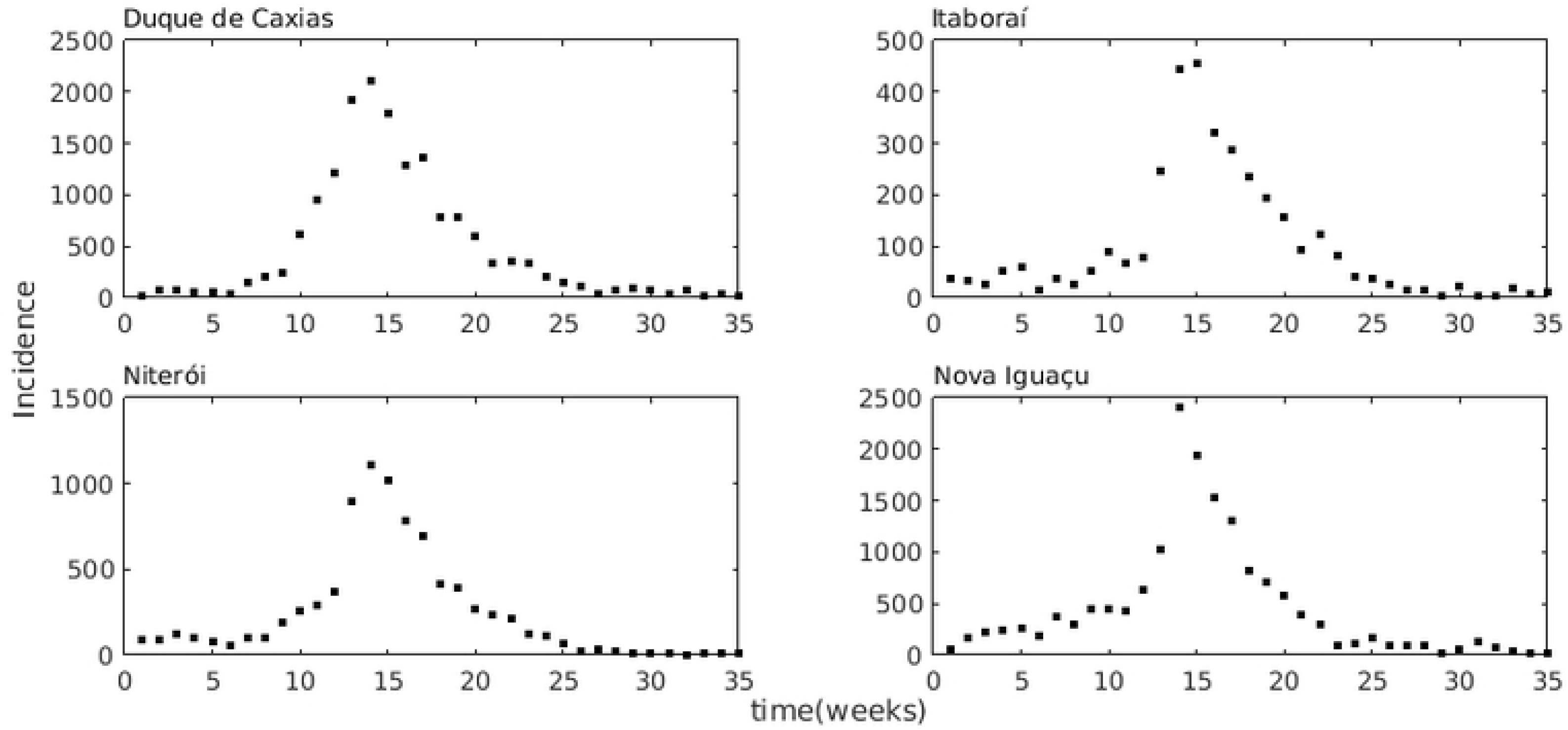
Incidence of dengue. Incidence of dengue in Duque de Caxias, Itaboraí, Niterói and Nova Iguaçu in the period from the 1st week of 2008 to the 35th week of 2008.

**Table 1.** Parameters that are fixed in all simulations; the total population of each city *N_h_i__* and the human birth/mortality rate *μ_h_*.

| Cities | $N_{h_i}$ | $\mu_h$ (weeks) |
| --- | --- | --- |
| Duque de Caxias | 855048 | $1/(70*52)$ |
| Itaboraí | 218008 | $1/(70*52)$ |
| Niterói | 487562 | $1/(70*52)$ |
| Nova Iguaçu | 796257 | $1/(70*52)$ |

## Results

Regarding the initial values of the parameters, we will consider that the initial susceptible population is 90% of the total population of each city due to the fact that dengue is endemic in Brazil, so *S_r_* (0) = 0.90 × *N_hr_*. The human recovery rate is initially *γ* =1 (week). Also the parameters of mobility start with the values *d_rs_* = 0.0001, being able to assume values in the interval [0, 1], where the indices represent the proportion of the population that leaves City *r* and goes to City *s*.

Recalling that by definition: 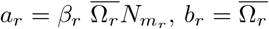 and 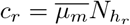. As a consequence of considering the System (11) with time scale separation, we have that the mosquitoes equations do not appear in the system, however the values *a_r_, b_r_* and *c_r_, r* =1, 2 depend on the mosquito’s parameter, being necessary to make an initial assumption for its values. Lets consider that initially, *μ_m_* = 7/10 (weeks), so we use an initial approximation for *β_r_* as *β_r_* = 2*γ*, Ω_*r*_ as Ω_*r*_ = 1 *μ_m_* and *N_m_r__* = *mN_h_r__*. Actually, by defining the parameter *ε* = *μ_h_*/*μ_m_*, it follows that

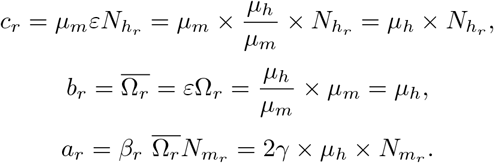

Thus, we have that the total mosquito population *N_m_r__*, is the only parameter of the mosquitoes being used, which interferes with the initial value of the parameter *a_r_*. Therefore, we will analyze the fit of the model to the data considering: *N_m_r__* = *N_h_r__, N_m_r__* = 2*N_h_r__, N_m_r__* = 3*N_h_r__* and *N_m_r__* = 4*N_h_r__*, and also by varying the initial amount of infected individuals *I_r_* = 1, 2, 3, 4. We will present the results with the value of *N_m_r__* that best fits the data, that is, the one that results in smaller quadratic error. Simulations with the mosquitoes population larger than 4 times the human population were not successful.

The results of the estimated parameters are presented in Table 2 for each pair of cities, and in the sequence the figures with their respective adjustments, where the incidence is the number of new cases recorded per week (black dots) and the solid lines in red are the amount of infected individuals obtained with the model according to the estimate obtained.

**Table 2.**
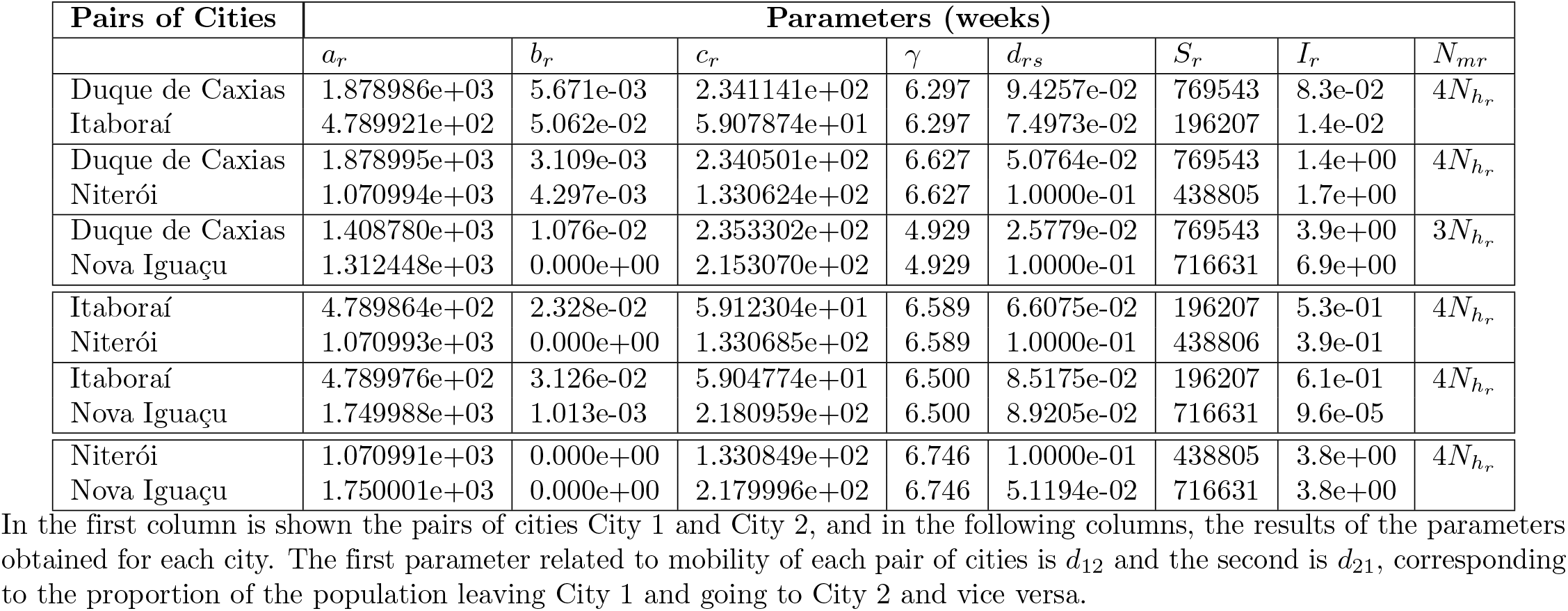
Results of parameters estimation for all pairs of cities.

**Fig 2.**
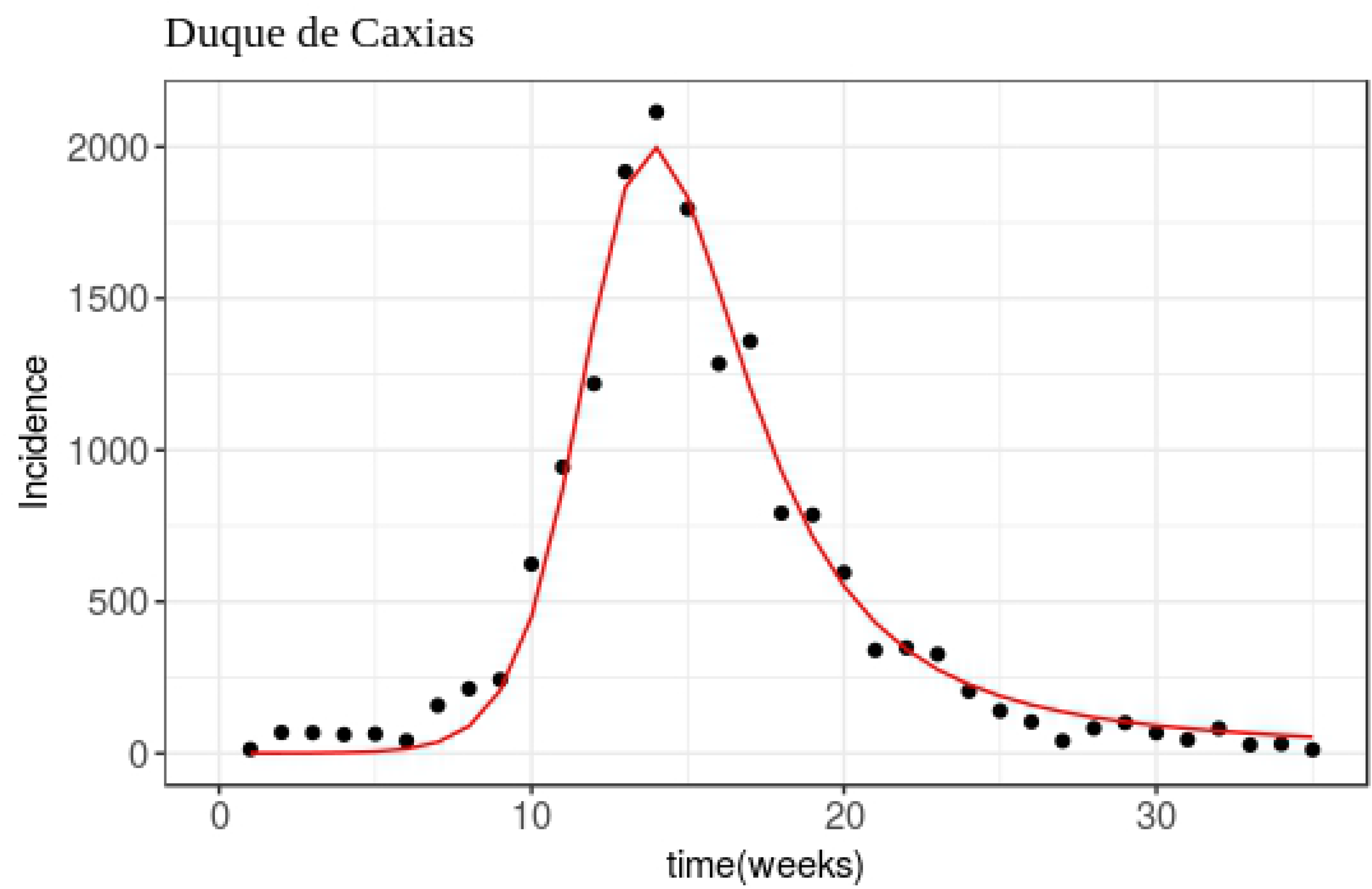
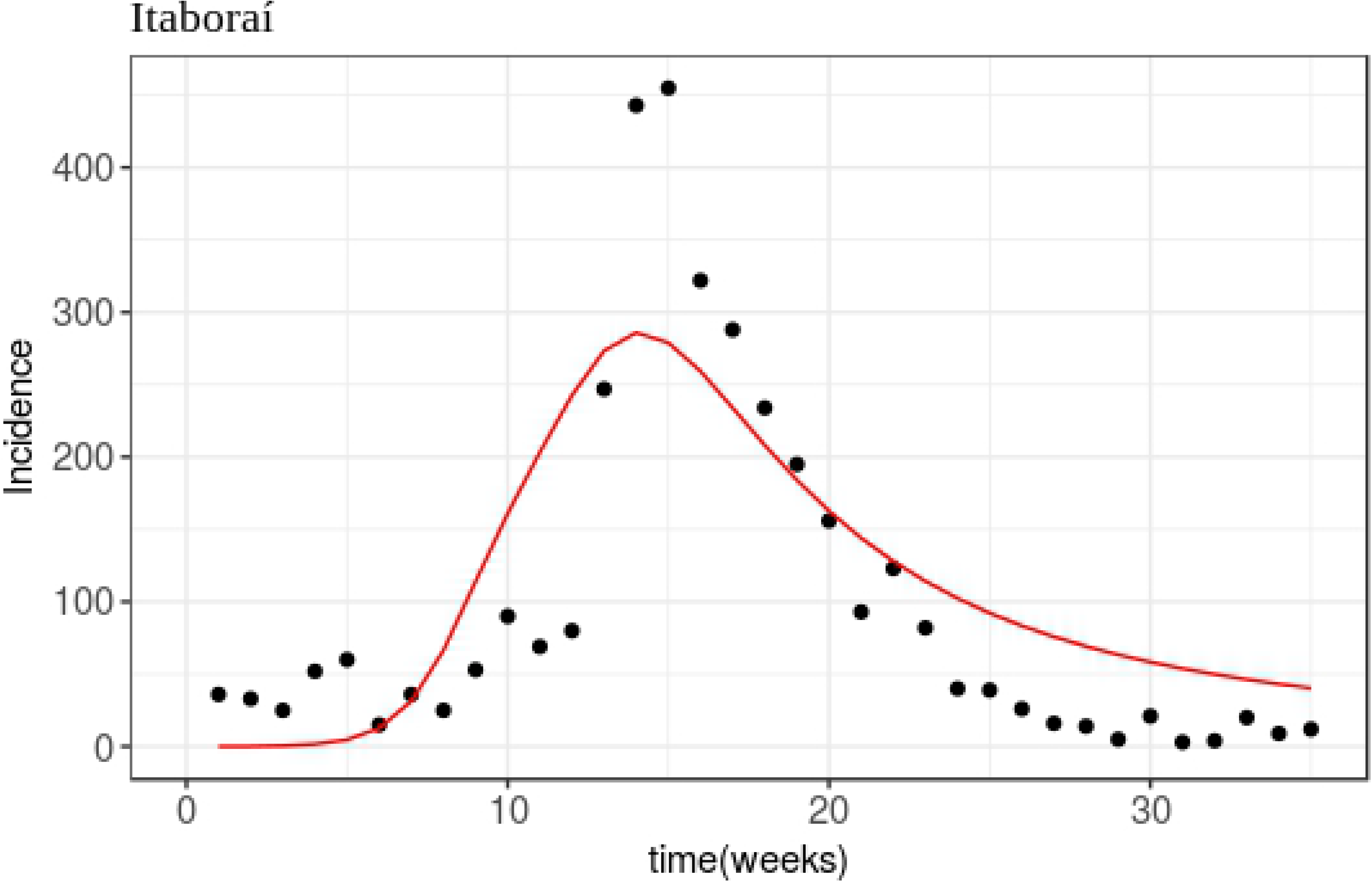
Fitting for Duque de Caxias and Itaboraí. Result of adjusting the Model 11 to dengue data of the cities: Duque de Caxias and Itaboraí, according to the parameters of Table 2.

**Fig 3.**
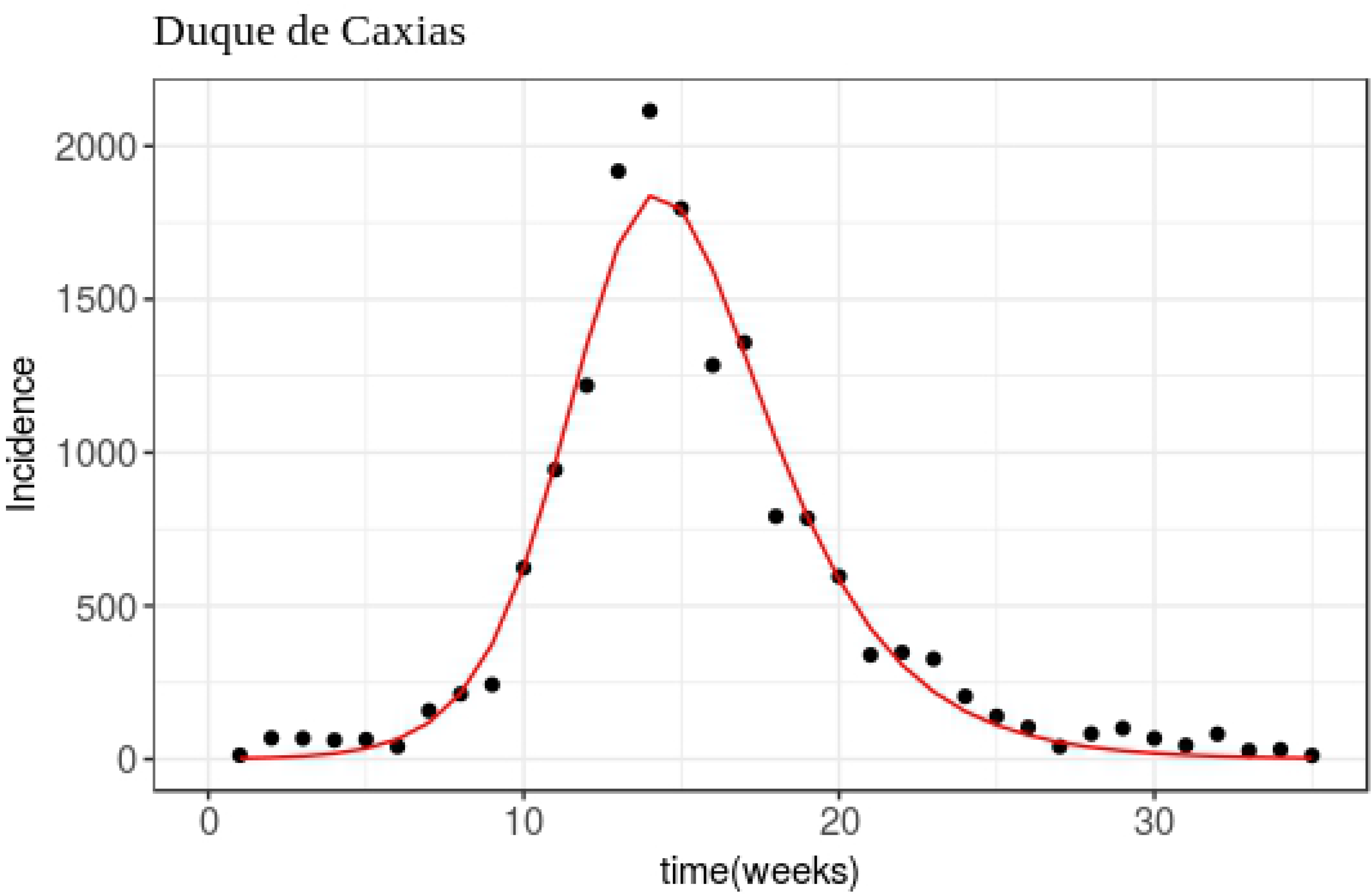
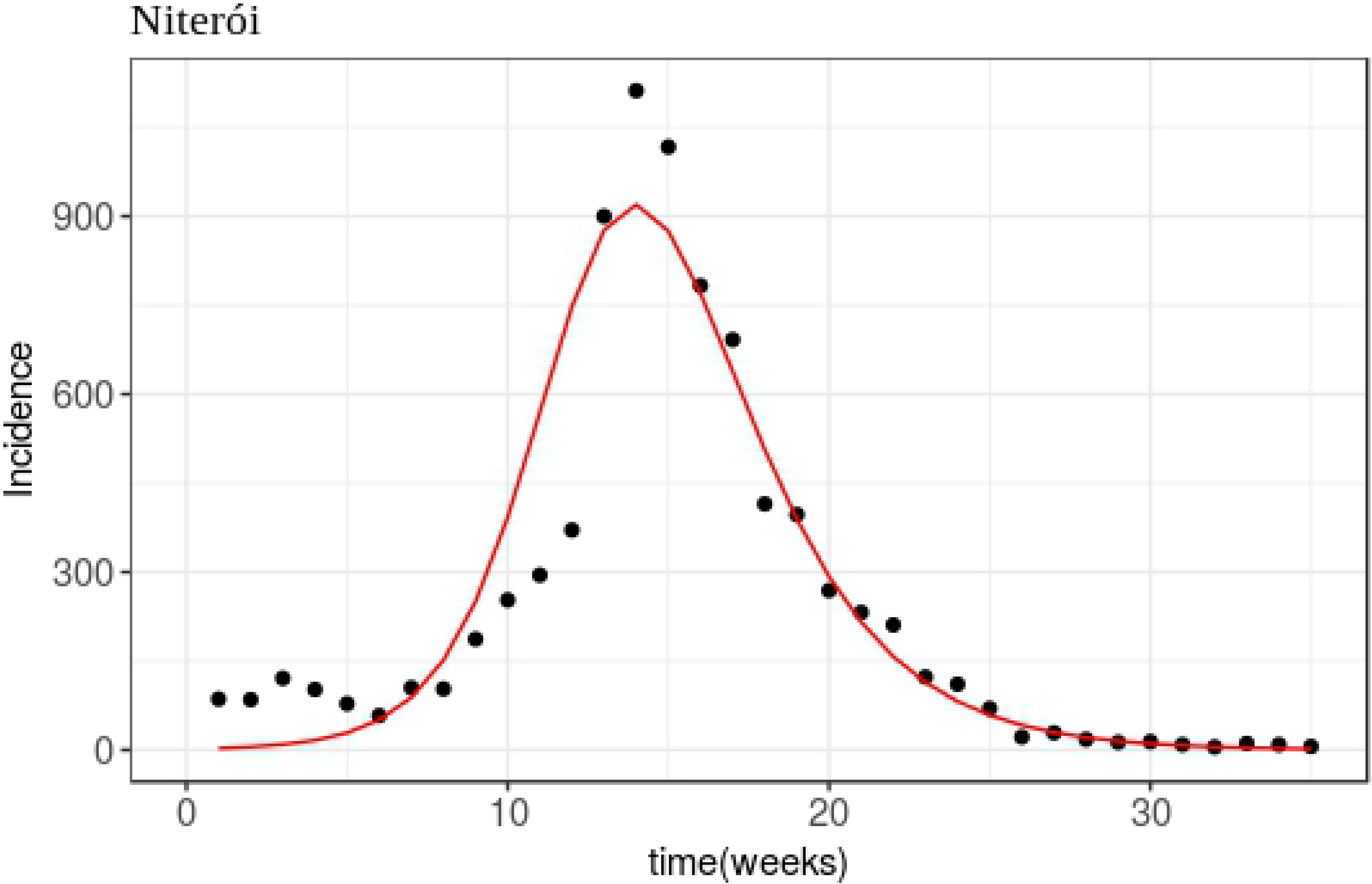
Fitting for Duque de Caxias and Niterói. Result of adjusting the Model 11 to dengue data of the cities: Duque de Caxias and Niterói, according to the parameters of Table 2.

**Fig 4.**
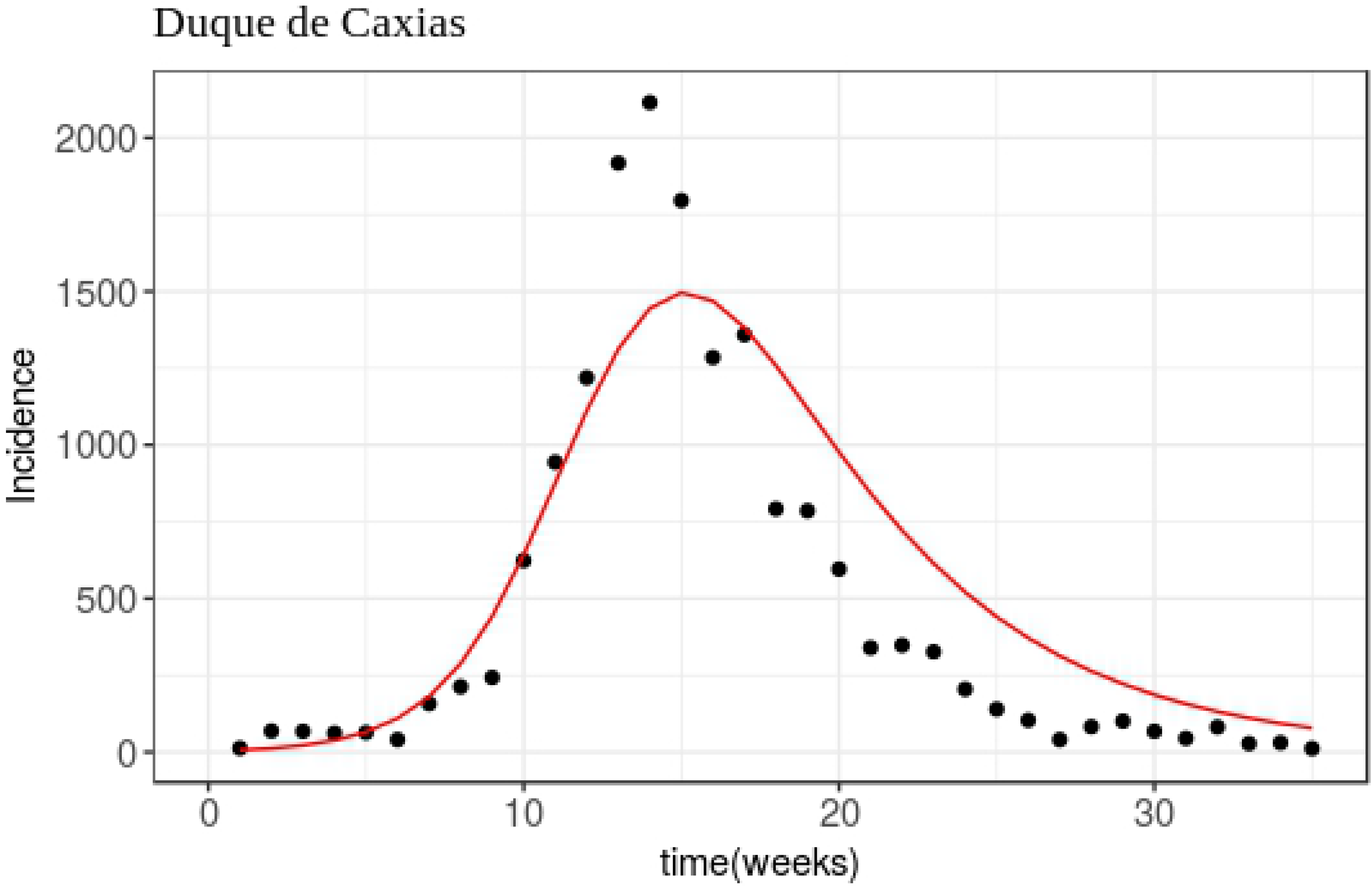
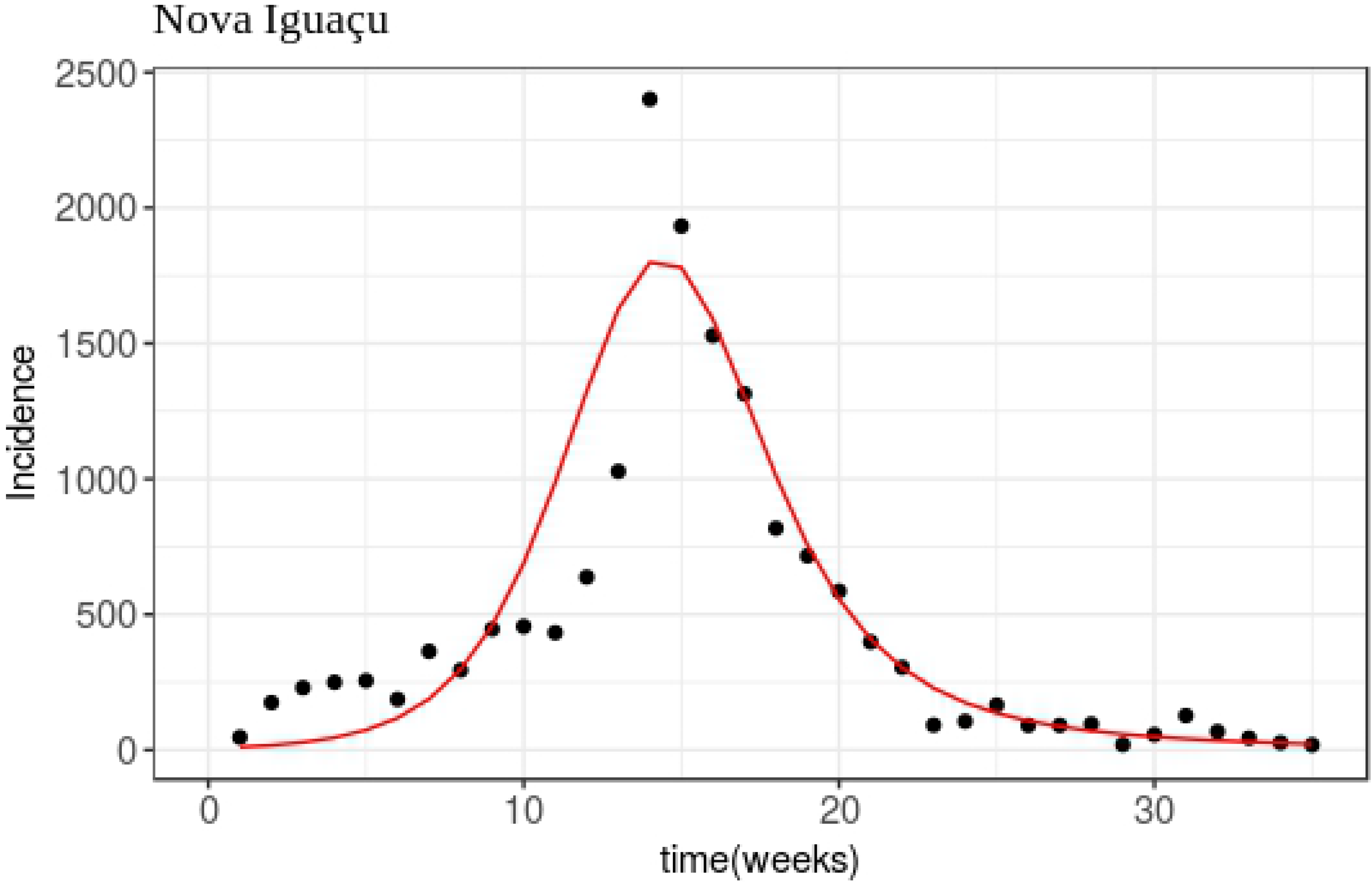
Fitting for Duque de Caxias and Nova Iguacu. Result of adjusting the Model 11 to dengue data of the cities: Duque de Caxias and Nova Iguacu, according to the parameters of Table 2.

**Fig 5.**
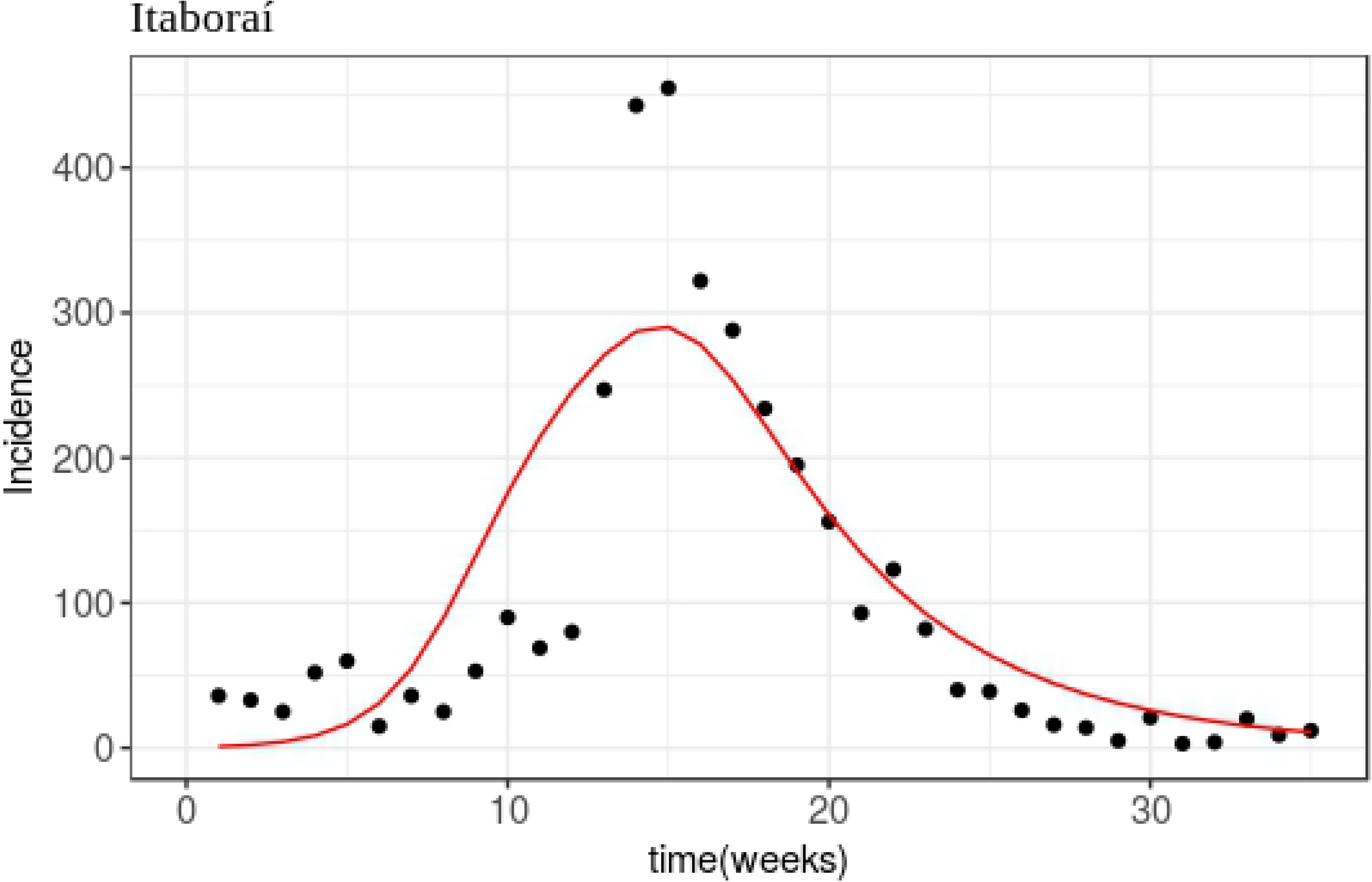
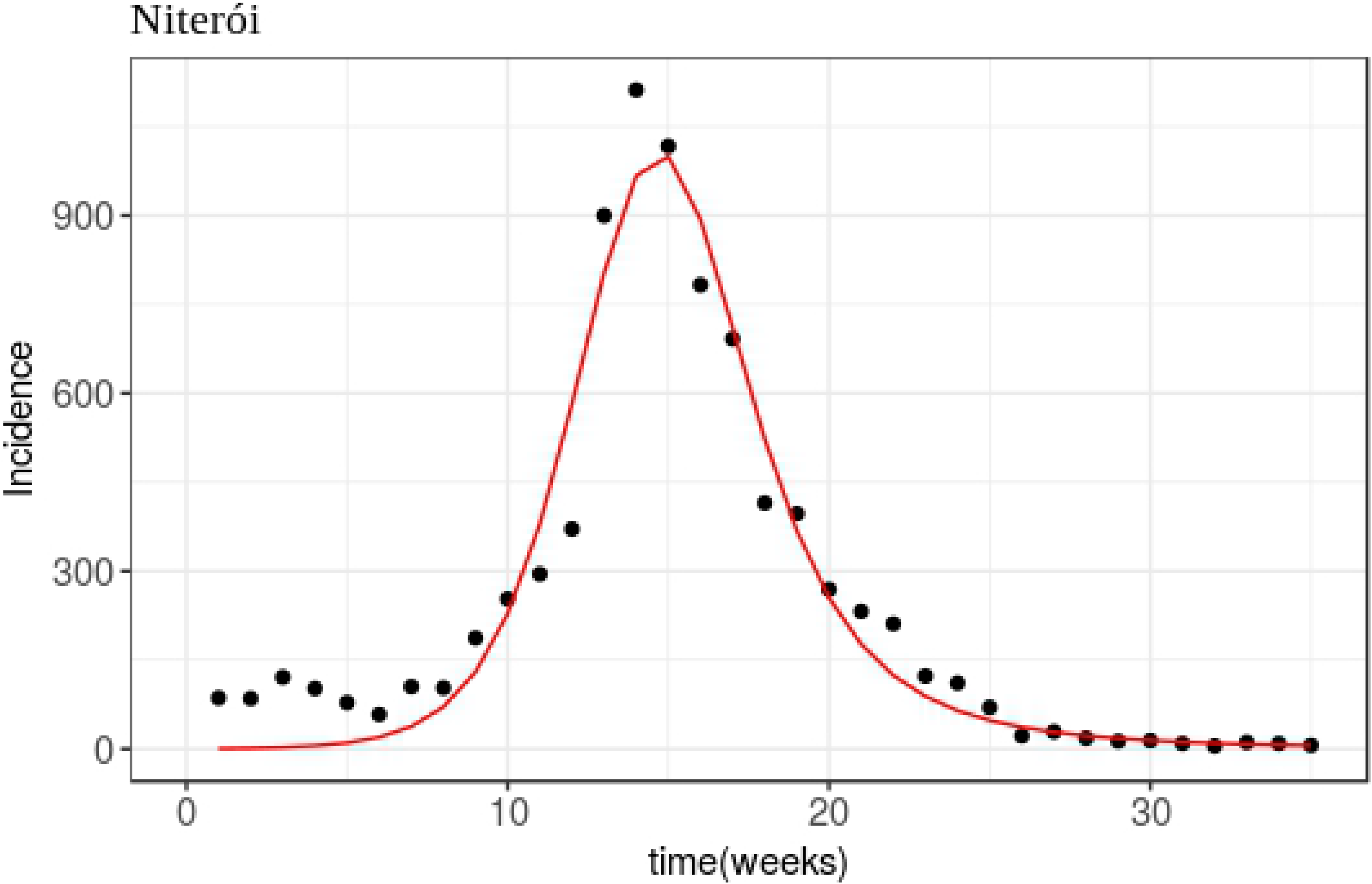
Fitting for Itaboraí and Niterói. Result of adjusting the Model 11 to dengue data of the cities: Itaboraí and Niterói, according to the parameters of Table 2.

**Fig 6.**
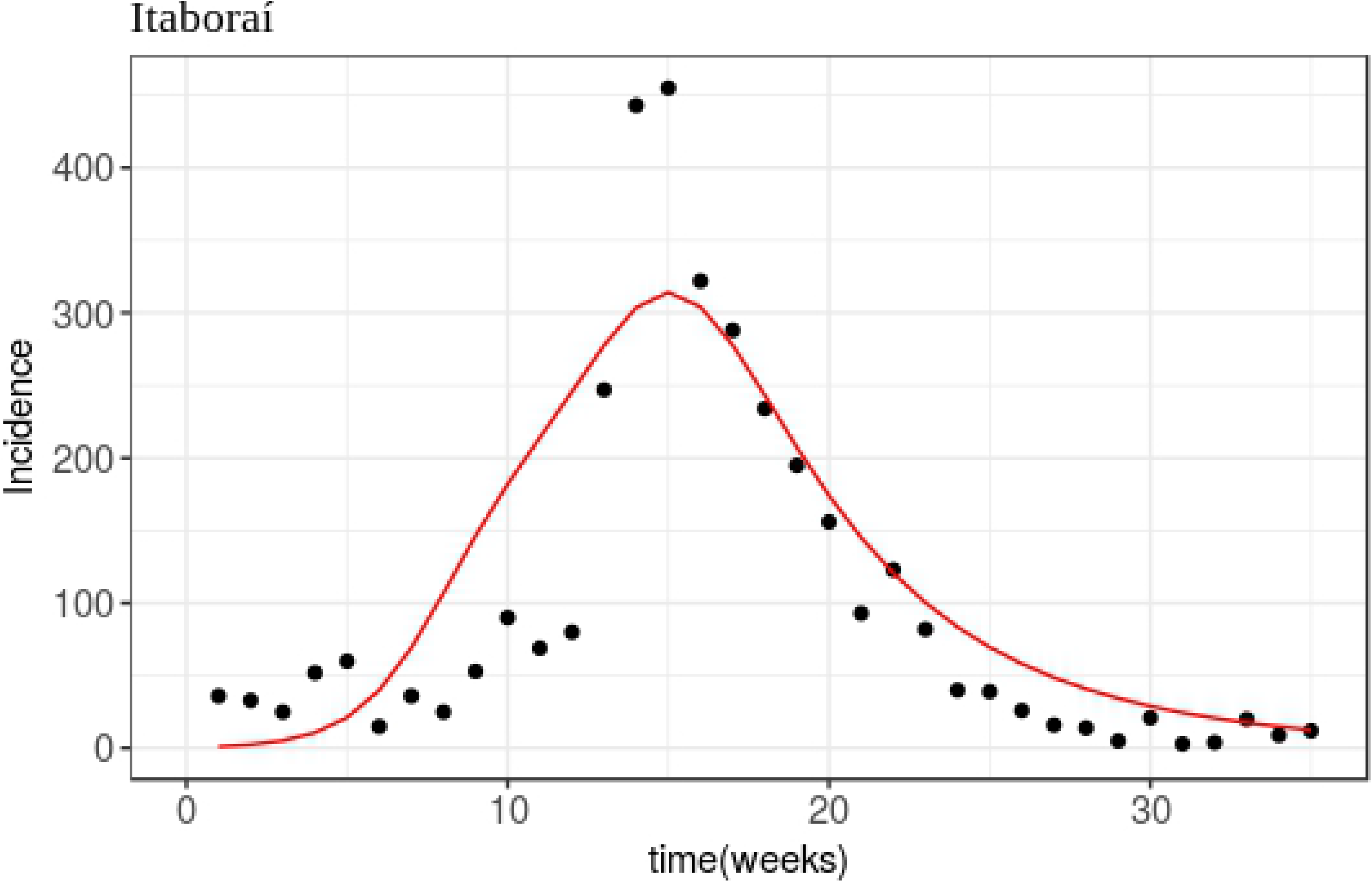
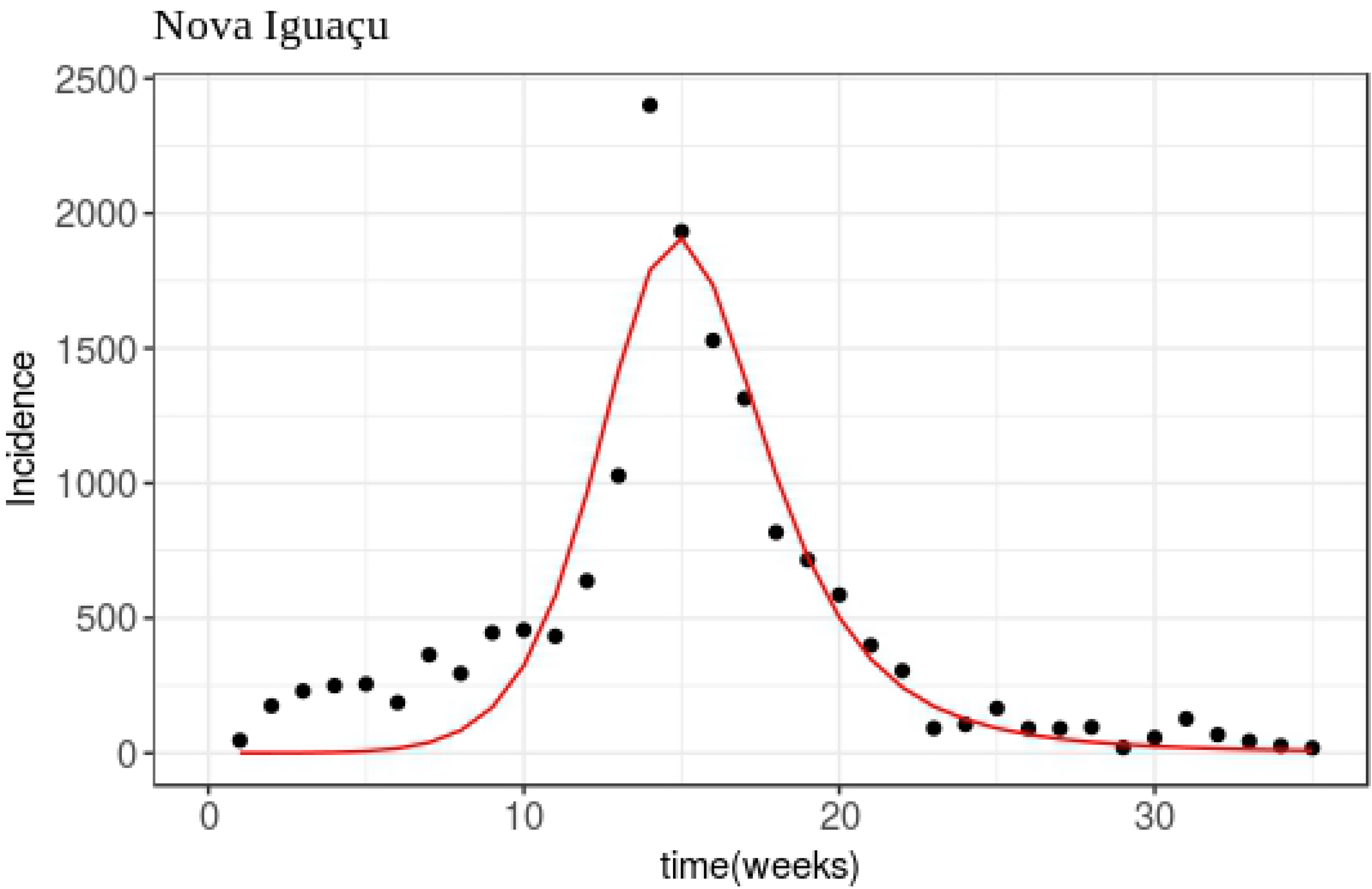
Fitting for Itaboraí and Nova Iguacu. Result of adjusting the Model 11 to dengue data of the cities: Fitting for Itaboraí and Nova Iguaçu, according to the parameters of Table 2.

**Fig 7.**
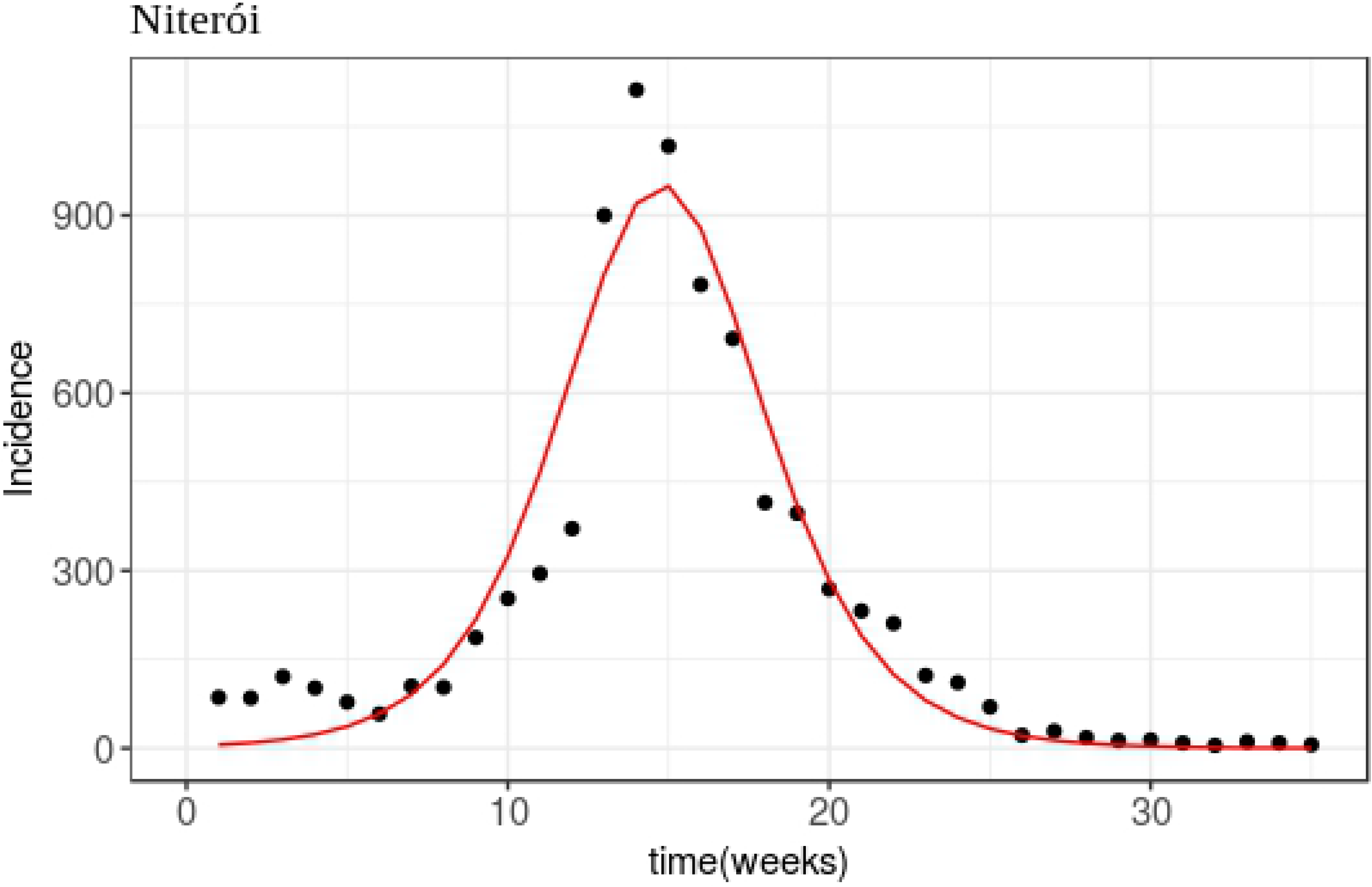
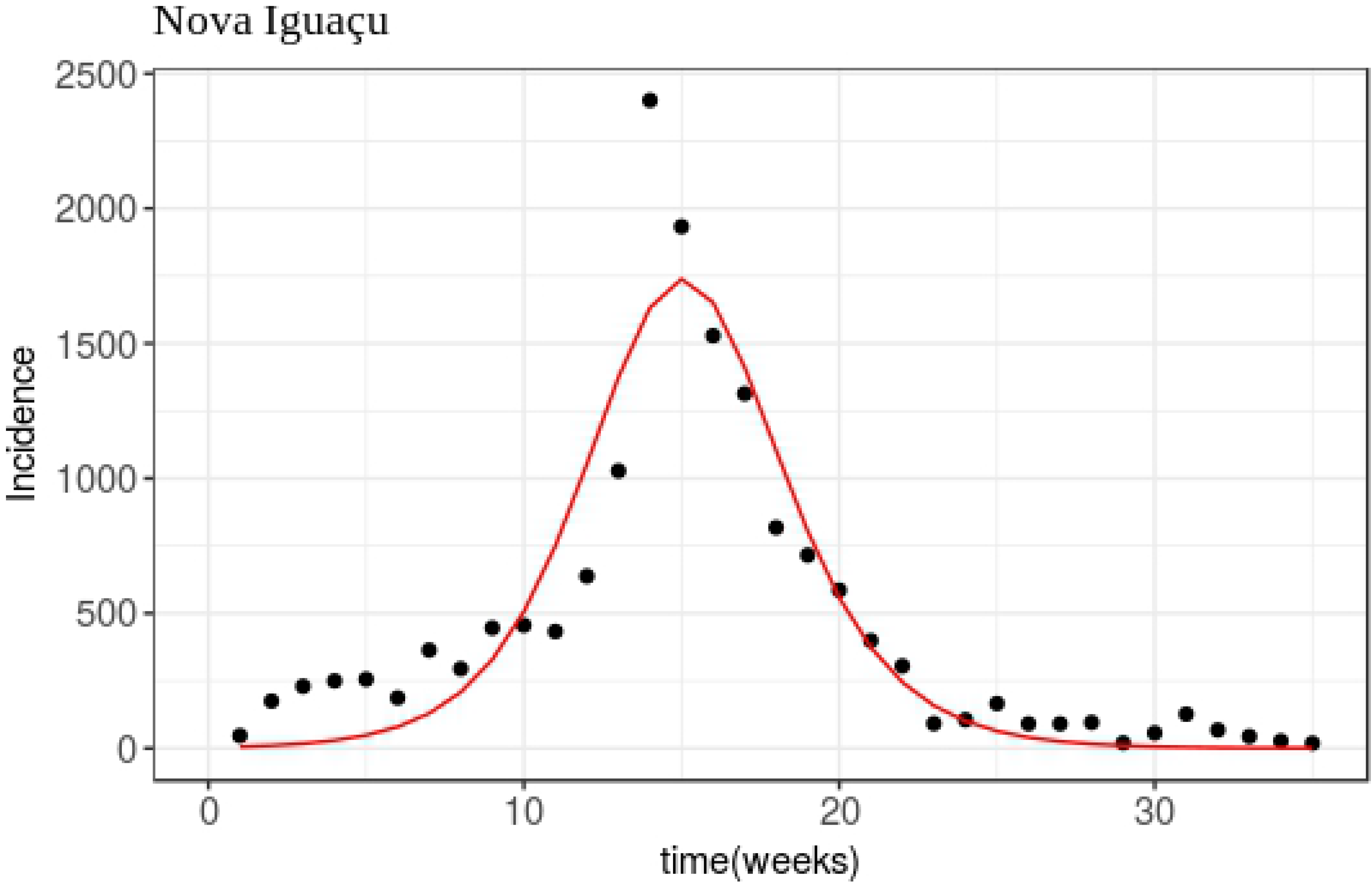
Fitting for Niteroi and Nova Iguacu. Result of adjusting the Model 11 to dengue data of the cities: Niterói and Nova Iguaçu, according to the parameters of Table 2.

## Discussion

From a *SIRS_m_I_m_* model with humans and vectors, we made a separation of time scales and reduced the system to an equivalent system independent of the mosquitoes equations, and then we add mobility in this new system. So, the new parameters depend only on the respective parameters for humans and the total mosquitoes population.

We made some considerations about the initial value of the parameters, and then we applied an algorithm to fit the model to dengue data of some cities in Rio de Janeiro state. The results showed that the reduced model was able to successfully adjust the beginning of the dengue outbreak for all pairs of cities, also obtaining values for the parameters related to mobility. This gives us an indication that human mobility actually has influence on the spread of dengue.

In addition, we have that the model with time-scale separation *S_r_I_r_* depends only on one parameter related to mosquitoes, whereas the *SIRS_m_I_m_* model contains three parameters of the vectors, which makes it more difficult to fit the model to the data due to the lack of available information about mosquitoes, since the time series of dengue provide only the amount of humans infected weekly. This gives us perspectives for a future study involving more complex mobility networks applied to analyse vector-borne diseases.

## Acknowledgments

The second author (MCP) is partially supported by CNPq 303253/2017-7 and FAPESP 2017/02630-2 (Brazil).

